# α7 nicotinic acetylcholine receptor signaling modulates the inflammatory and iron homeostasis in fetal brain microglia

**DOI:** 10.1101/097295

**Authors:** M. Cortes, M. Cao, H.L. Liu, C.S. Moore, L.D. Durosier, P. Burns, G. Fecteau, A. Desrochers, L.B. Barreiro, J.P. Antel, M.G. Frasch

**Affiliations:** Dept. of Obstetrics and Gynaecology and Dept. of Neurosciences, CHU Ste-Justine Research Centre, Faculty of Medicine, Montréal, QC, Canada; Animal Reproduction Research Centre (CRRA), Faculty of Veterinary Medicine, Université de Montréal, Montréal, QC, Canada; Neuroimmunology Unit, Montréal Neurological Institute, McGill University, Montréal, QC, Canada; Dept. of Clinical Sciences, Faculty of Veterinary Medicine, Université de Montréal, QC, Canada; Dept. of Pediatrics, CHU Ste-Justine Research Centre, Faculty of Medicine, Université de Montréal, Montréal, QC, Canada; Dept. of Obstetrics and Gynaecology, University of Washington, Seattle, WA, USA

**Author notes:** Address of correspondence: Martin G. Frasch, Department of Obstetrics and Gynecology, University of Washington, 1959 NE Pacific St, Box 356460, Seattle, WA 98195. M. Cortes and M. Cao contributed equally to this manuscript.

## Abstract

Neuroinflammation *in utero* may result in life-long neurological disabilities. Microglia play a pivotal role, but the mechanisms are poorly understood. No early postnatal treatment strategies exist to enhance neuroprotective potential of microglia. We hypothesized that agonism on α7 nicotinic acetylcholine receptor (α7nAChR) in fetal microglia will augment their neuroprotective transcriptome profile, while the antagonistic stimulation of α7nAChR will achieve the opposite. Using an *in vivo* - *in vitro* model of developmental programming of neuroinflammation induced by lipopolysaccharide (LPS), we validated this hypothesis in primary fetal sheep microglia cultures re-exposed to LPS in presence of a selective α7nAChR agonist or antagonist. Our RNAseq and protein level findings show that a pro-inflammatory microglial phenotype acquired *in vitro* by LPS stimulation is reversed with α7nAChR agonistic stimulation. Conversely, antagonistic α7nAChR stimulation potentiates the pro-inflammatory microglial phenotype. Surprisingly, under conditions of LPS double-hit an interference of a postulated α7nAChR - ferroportin signaling pathway may impede this mechanism. These results suggest a therapeutic potential of α7nAChR agonists in early re-programming of microglia in neonates exposed to *in utero* inflammation via an endogenous cerebral cholinergic anti-inflammatory pathway. Future studies will assess the role of interactions between inflammation-triggered microglial iron sequestering and α7nAChR signaling in neurodevelopment.

## INTRODUCTION

Brain injury acquired antenatally remains a major cause of long-term neurodevelopmental sequelae in children and adults.(1) Although the etiology of antenatal brain injury is undoubtedly multifactorial, there is growing evidence for a role for maternal and fetal infection and inflammation(2),(3),(4), which is supported by animal studies.(5),(2),(6) Both systemic and neuroinflammation have been implied as important pathophysiological mechanisms acting independently to cause fetal brain injury or contributing to *in utero* asphyxial brain injury with consequences for postnatal health.(7),(8)

The main cause of fetal inflammation, chorioamnionitis, is a frequent (10% of all pregnancies, up to 40% of preterm births) and often subclinical fetal inflammation associated with ˜9fold increased risk for cerebral palsy spectrum disorders with life lasting neurological deficits and an increased risk for acute or life-long morbidity and mortality, inversely correlated with gestational age at delivery.(8),(9),(10)

In addition to short-term brain damage, recent studies suggest that neuroimmune responses to *in utero* infection may also have long-term health consequences. In adults, exposure to inflammatory stimuli can activate microglia (glial priming, reviewed in(11),(12)). Confronted with a renewed inflammatory stimulus, they can sustain chronic or exaggerated production of pro-inflammatory cytokines associated with postnatal neuroinflammatory diseases such as Multiple Sclerosis or sustained cognitive dysfunction (“second hit” hypothesis).(12),(13),(14)

α7 nicotinic acetylcholine receptor (α7nAChR) signaling in microglia may be involved in modulating TNF-α release to push microglia towards a neuroprotective role under conditions of lipopolysaccharide (LPS) exposure.(15),(16),(17)

We hypothesized that agonistic stimulation of α7nAChR in fetal microglia will augment their neuroprotective transcriptome profile, while the antagonistic stimulation of α7nAChR will achieve the opposite. Using a novel *in vivo* - *in vitro* model of developmental programming of neuroinflammation induced by LPS, we validate this hypothesis in primary fetal sheep microglia cultures exposed to LPS in presence of a selective α7nAChR agonist or antagonist.

## RESULTS

### Primary fetal sheep microglia culture

*In vitro* studies were conducted in primary cultures derived from six controls (naïve) and from two *in vivo* LPS-exposed animals (SH) in 1-2 *in vitro* replicates each animal depending on cell numbers obtained (Figure 1A). First, we investigated cytokine secretion properties of microglial cultures in the absence or presence of LPS. Methodology and results are presented elsewhere.(18) Second, we studied IL-1β secretion profile (Figure 1B) in response to LPS accompanied by co-incubation with α7nAChR agonist AR-R17779 (NA or SHA) and the α7nAChR antagonist α-Bungarotoxin (NB or SHB).

**Figure 1.**
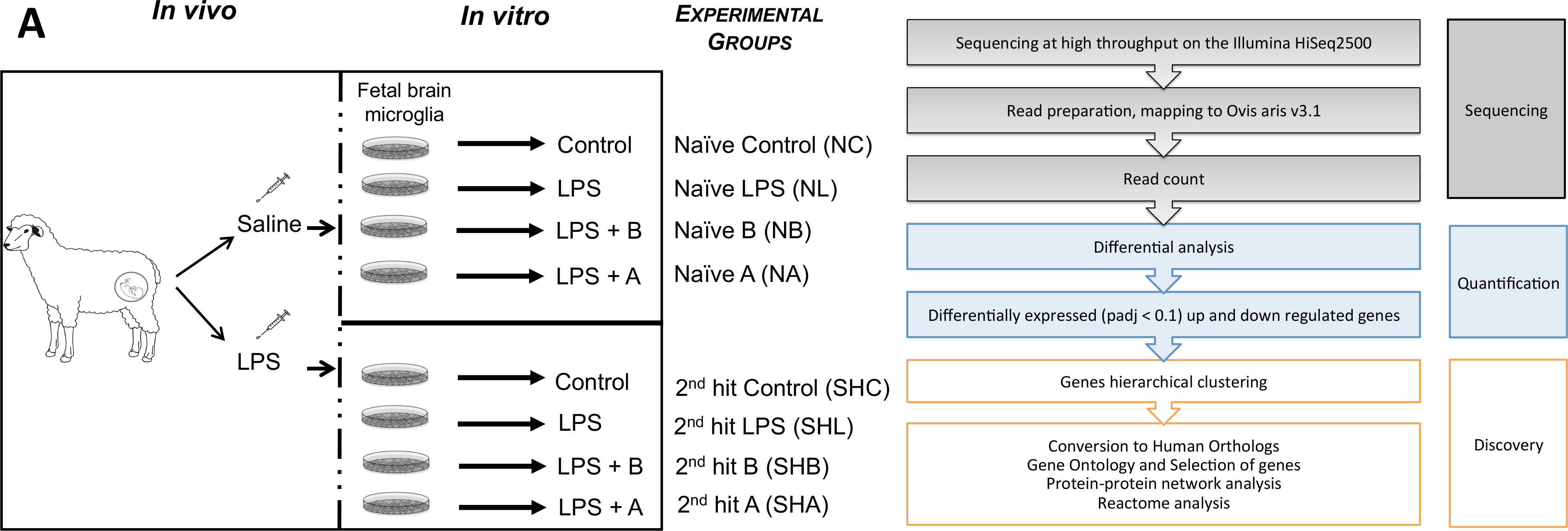
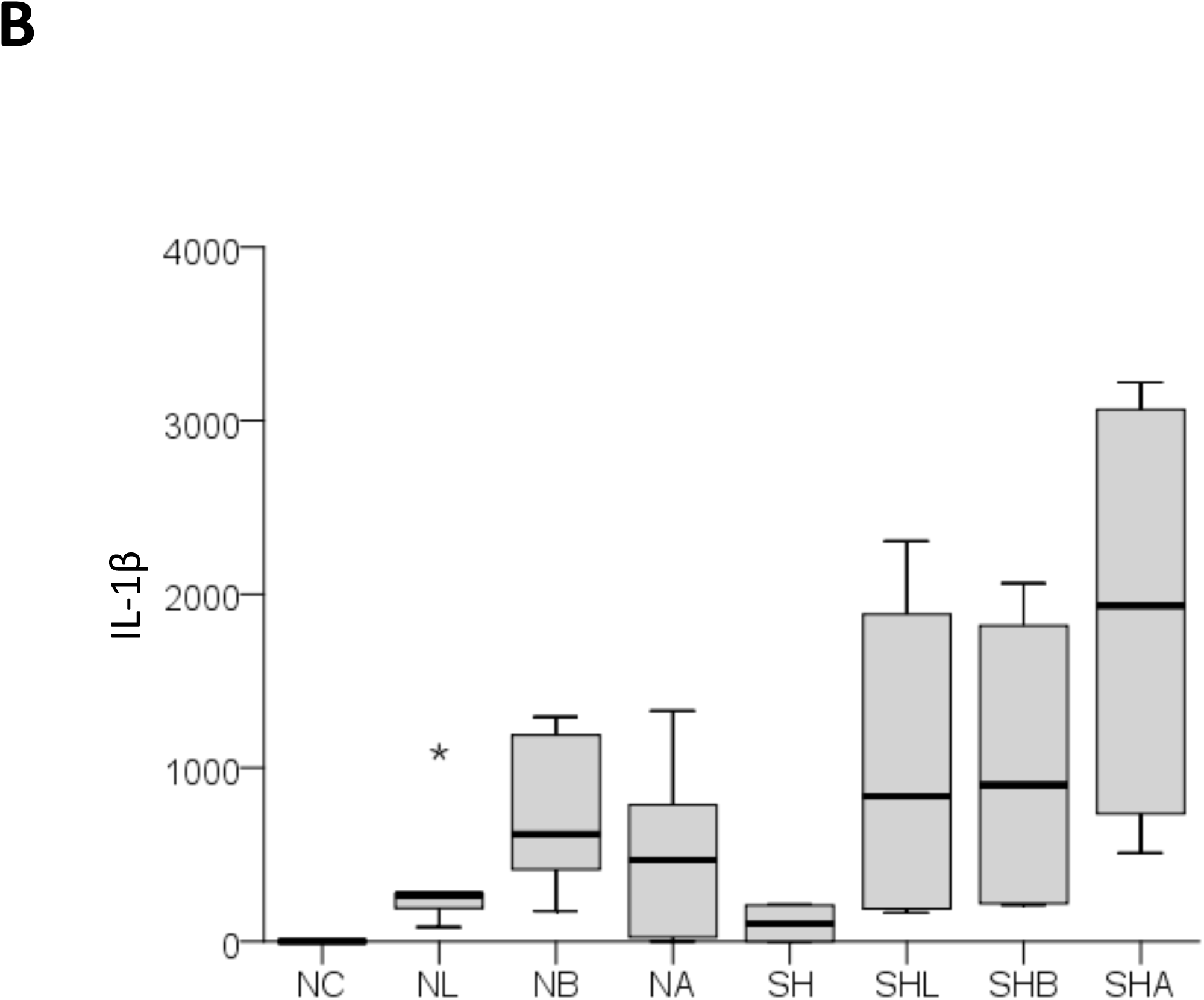
Experimental design of modulation through α7nAChR signaling in a double-hit fetal sheep model. **A.** *In vivo, in vitro* and RNAseq experiments are illustrated. *In vivo* study, Control (saline); *in vitro* study, cultured cells derived from *in vivo* Control animal, named as Naïve, there are 8 experimental groups: naïve Control (NC), naïve LPS (NL), naïve exposed to α-Bungarotoxin (NB), naïve exposed to AR-R17779 (NA), and each respective second-hit groups (SH). For RNAseq data comparisons, the group SHA was excluded. **B.** Box plot graph of cytokine IL-1ß response obtained for each group of replicates. * represents an outlier outside 95^th^ percentile. GEE model results are presented in text and no significance marks are provided in the figure. Briefly, we found significant main and interaction term effects (p<0.05) for LPS and drug treatment and the contribution of *in vivo* LPS exposure, *i.e.*, SH effect on IL-1β secretion profile.

The cytokine IL-1β secretion profile showed a non-random distribution pattern (p<0.001, main term “group”), but not for the main term “SH yes or no” (p=0.122). The latter main term identified each group as having or not having been exposed to an *in vivo* hit, *i.e.*, testing for the SH effect on IL-1β secretion profile <u>without</u> accounting for the experimental group. LPS exposure led to IL-1β rise (p=0.020) which was non-randomly changed by α7nAChR agonism (p<0.001), but not by antagonism (p=0.801).

The GEE model exploring the contribution of the second hit to the IL-1β secretion profile revealed a significant interaction between the four experimental groups (control, LPS, antagonist and agonist pre-treatments) and the presence or absence of two hits (p<0.001 for interaction term “group”*”SH yes or no”). Specifically, <u>without</u> the preceding *in vivo* hit, α7nAChR <u>a</u>gonism reduced the effect of this heightened IL-1β secretion (p=0.028). Surprisingly, <u>with</u> the preceding *in vivo* hit, *i.e.*, in the SH groups, α7nAChR agonism amplified the effect (p=0.028). That is, *in vivo* exposure to LPS reversed the effect of the agonistic α7nAChR stimulation. Meanwhile, for α7nAChR <u>anta</u>gonism the results were consistently supporting the initial hypothesis with IL-1β rising when accounting in the model for the preceding absence or presence of the *in vivo* first LPS hit (interaction terms “NB”* “SH yes or no” (p=0.048). We found no significant interaction for NC and NL groups and SH (p=0.204).

### RNAseq approach

#### Whole transcriptome analysis

We reported the genomic landscape of primary fetal sheep microglia in response to LPS using similar quality control methods to confirm the cell culture purity.(18) Here we sequenced at high throughput the whole transcriptome of microglia exposed to LPS and pre-incubated with α7nAChR agonist or antagonist (Figure 1A). We performed a direct differential analysis of NA versus NB which eliminated the background noise of NC. This approach allowed us to observe the immediate effect of LPS on microglial transcriptome when it is modulated by α7nAChR antagonist versus agonist treatments. We performed 6 differential analyses of microglia exposed to agonist and antagonist (NA, NB, respectively) versus control (NC) and second hit microglia exposed to antagonistic treatment (SHB). In Table 1, we summarized the number of differentially expressed genes (DEG) found for each differential analysis. Overall, the microglial transcriptome exposed to agonistic and antagonistic drugs revealed a greater amount of DEG than LPS-exposed microglia (latter results were published).(18)

**Table 1.** Differential analysis summary in naïve and second hit microglia after modulation of α7nAChR signaling. Differential analysis of count data was done with the DESeq2 package. Differential expressed genes were selected for padj < 0.1. Up regulation and down regulation represent positive and negative Log2 fold changes, respectively.

|  | NC - NB | NC - SHB | NB - SHB | NC - NA | NL - NA | NB-NA |
| --- | --- | --- | --- | --- | --- | --- |
| DE genes* | 2,400 | 7,314 | 7,340 | 2,007 | 2 | 162 |
| DE* up regulated | 1,432 | 4,351 | 4,086 | 1,103 | 2 | 24 |
| DE* down regulated | 968 | 2,963 | 3,254 | 904 | 0 | 138 |

\* padj < 0.1

**Table 2.**
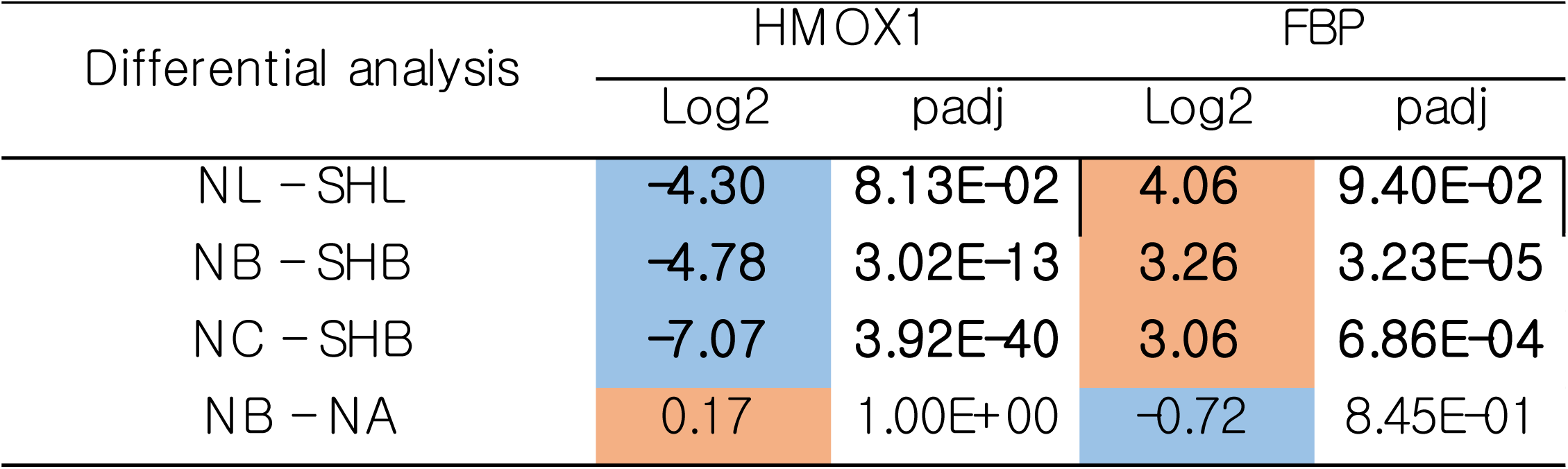
Genetic expression of HMOX1 and FBP after exposure to LPS and signaling through α7nAChR. Confirming our previous findings and the notion of the pro-inflammatory effect of blocking signaling through α7nAChR, we found HMOX1 to be progressively stronger down regulated and FBP up regulated due to a double-hit LPS exposure and subsequent pre-treatment with the receptor antagonist (SHB).

| Differential analysis | HMOX1 |  | FBP |  |
| --- | --- | --- | --- | --- |
|  | Log2 | padj | Log2 | padj |
| NL – SHL | -4.30 | 8.13E-02 | 4.06 | 9.40E-02 |
| NB – SHB | -4.78 | 3.02E-13 | 3.26 | 3.23E-05 |
| NC – SHB | -7.07 | 3.92E-40 | 3.06 | 6.86E-04 |
| NB – NA | 0.17 | 1.00E+00 | -0.72 | 8.45E-01 |

#### Unique transcriptome signature of agonistic and antagonistic stimulation in microglia

We identified 1,432 DEG (padj < 0.1) up regulated genes in NB microglia compared to NL. We compared the population of DE up regulated genes in NB with those in NL microglia: 1,234 genes were unique to NB. The analysis of pathways with Toppcluster revealed that unique genes to NB are members of the Jak-STAT, TNF-α and NFKB signaling pathways (Table 1, Figure 2A). 968 DE down regulated genes were identified in NB microglia compared to NL. All 53 genes previously identified in NL were also DE down regulated in NB. Thus, NB showed a unique signature of 915 genes. Gene ontology of genes unique to NB revealed that these genes are mostly part of coenzyme binding (GO:0050662 and P = 8.37 × 10^−6^), GTPase regulator activity (GO:0030695 and P = 3.11 × 10^−5^), damaged DNA binding activity (GO:0003684 and P = 2.59 × 10^−4^) and H4 histone acetyltransferase activity (GO:0010485 and P = 9.42 × 10^−4^).

**Figure 2.**
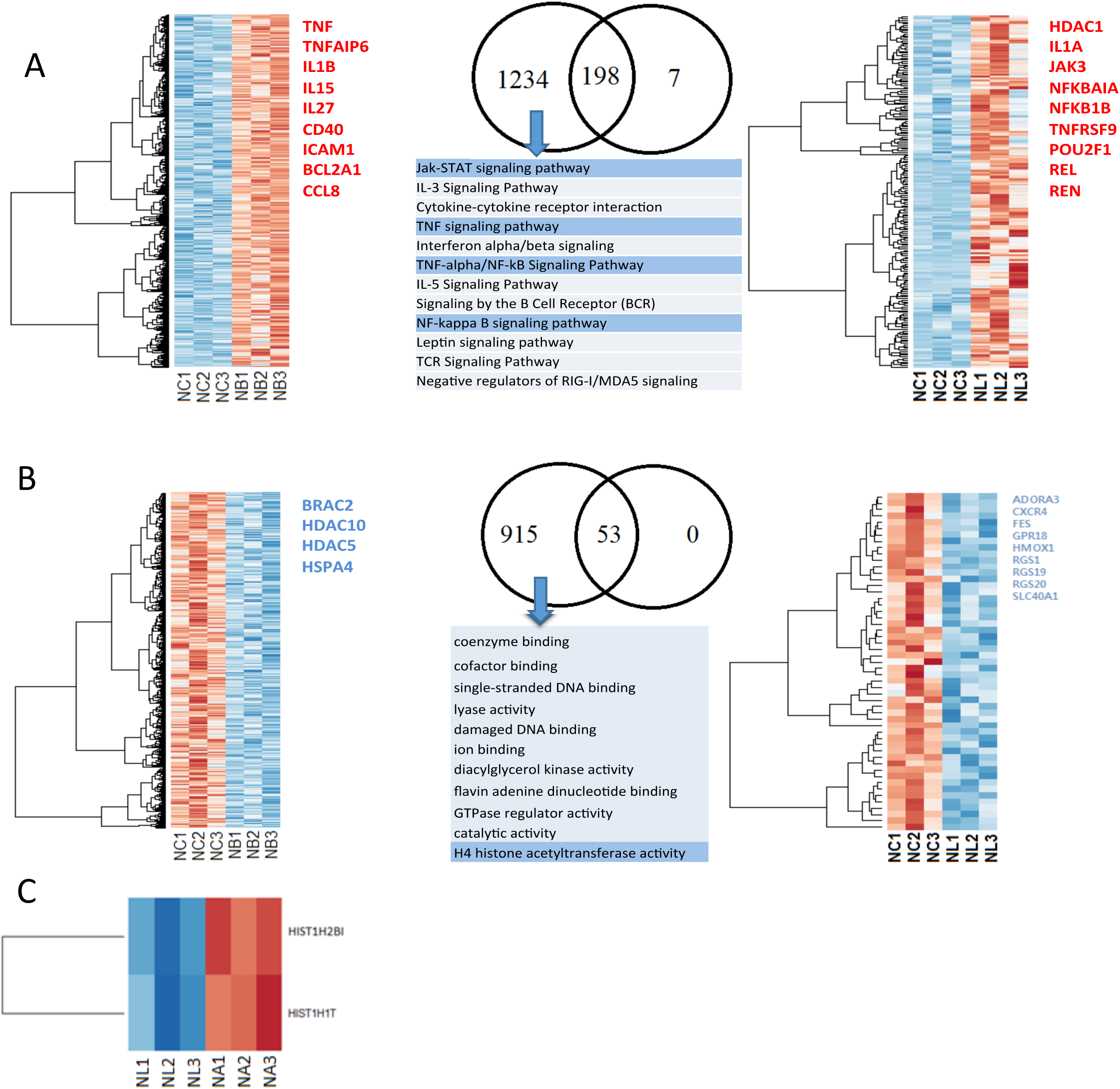
Stimulation of α7nAChR enhances the anti-inflammatory properties of fetal microglia. Differential analysis of the transcriptome of microglia pre-treated with the α7nAChR antagonist α-Bungarotoxin (NB) and the α7nAChR agonist AR-R17779 (NA) compared to controls (NC) and compared to the *in vitro* LPS-treated microglia whose α7nAChR activity was not modulated (NL). The Venn diagrams represent the number of unique genes for each group, and in the middle, the number of common genes. Selected genes of interest are written on the right side of each heat map. **A**. Microglia treated with the α7nAChR antagonist α-Bungarotoxin recruited more genes involved in the inflammatory pathway. **B**. α-Bungarotoxin also increased the response in DE down regulated genes. **C**. NL and NA microglia revealed two DE up regulated histone genes.

We extracted genes that were uniquely differentially expressed in NB and not found in LPS-exposed naïve microglia (NL vs. NC). We reported up regulation of genes involved in the NFKB and JAK-STAT pathway. (18) A closer look at all genes involved in these two pathways confirmed up regulation of TNF and IL1B. However, in our previous results, the latter two were not differentially expressed under LPS exposure alone. Our current differential analysis of microglia treated with α-Bungarotoxin prior to LPS exposure showed that both TNF and IL1B are differentially expressed and up regulated (padj = 1.17 × 10^−25^ and padj = 8.02 × 10^−88^, respectively), further confirming our hypothesis that antagonistic stimulation of α7nAChR potentiates LPS-triggered microglial inflammation (Figure 2A).

In a similar way, from both differential analyses to baseline NC, we extracted DE down regulated genes unique to NB and not found in NL. Here we found that HDAC10 and HDAC5 are uniquely DE down regulated in NB (padj = 8.81 × 10^−2^, padj = 5.24 × 10^−6^, respectively). Further analysis of HDAC genes is described below (Figure 2B, Table 3).

**Table 3.** Impact of α7nAChR signaling on memory of inflammation. Differential analysis of HDACs and HATs genes, DEG are indicated with a **bold** font (padj < 0.1). In our previous report, we highlighted the potential role of HDAC1, 2 and 6 in memory of inflammation in fetal microglia.

| Relevance | Gene | NC-NL | NC-SHC | NL-SHL | NB-NA | NC-NB | NC-NA | NB-SHB |
| --- | --- | --- | --- | --- | --- | --- | --- | --- |
| HDAC genes : Potential epigenetic regulators | HDAC1 | **2.271** | 0.145 | 0.676 | -0.533 | **3.006** | 2.470 | -0.690 |
|  | HDAC10 | -0.242 | -0.84 | 0.116 | 0.193 | **-1.109** | -0.914 | 1.140 |
|  | HDAC11 | -0.214 | 0.867 | 0.812 | 0.040 | -1.201 | -1.166 | 2.368 |
|  | HDAC2 | -0.299 | **-2.746** | -2.423 | 0.051 | 0.206 | 0.256 | **-2.719** |
|  | HDAC3 | 0.045 | -0.692 | 0.321 | -0.160 | 0.211 | 0.057 | -0.331 |
|  | HDAC4 | 1.292 | 1.502 | -0.691 | -0.515 | 0.761 | 0.243 | 0.718 |
|  | HDAC5 | -0.869 | 0.333 | 0.501 | 0.260 | **-1.614** | -1.355 | **0.889** |
|  | HDAC6 | **-0.688** | -0.126 | -0.43 | 0.129 | **-0.824** | -0.695 | 0.250 |
|  | HDAC7 | -0.109 | 0.643 | -0.486 | -0.534 | -0.156 | -0.690 | -0.474 |
|  | HDAC8 | 0.336 | -0.889 | -2.51 | -0.150 | **0.802** | 0.653 | **-2.623** |
|  | HDAC9 | 0.732 | 1.556 | -1.816 | -0.311 | -0.251 | -0.565 | 0.296 |
| Histone Acetyltransferase 1 | HAT1 | **-0.379** | -1.639 | -0.865 | 0.564 | -0.137 | 0.431 | **-1.470** |

When comparing NL to NA, we did not identify any DE down regulated genes. However, differential analysis of agonist-stimulated microglia rendered a unique signature compared to LPS-treated microglia. We found that two genes were DE up regulated in NA versus NL, ENS0ARG00000020076 and HIST1H1T (padj=1.01 × 10^−10^, padj = 4.98 × 10^−2^, respectively). Per ensemble database, the gene ENS0ARG00000020076 corresponds to the HIST1H2BI, a family member of the Histone cluster 1 H2B (Table 3, Figure 2C).

#### Effect of α7nAChR agonist and antagonistic drugs on microglial transcriptome

Our main analysis focused on differences between NB and NA treatment. Our differential analysis of NA versus NB revealed 162 DEG, among which 24 were upregulated and 138 were down regulated (Table 1). Gene ontology of DEG down regulated in NA versus NB showed that DE down regulated were associated with the immune system (GO:0002376), however two DE up regulated genes also clustered for the GO terms immune system, HSPA6 and GADD45G (Figure 3B). In the human genome, the gene HSPA6 codes for the Heat shock 70 kDa protein 6, and GADD45G codes for Growth arrest and DNA damage-inducible protein GADD45 gamma.

**Figure 3.**
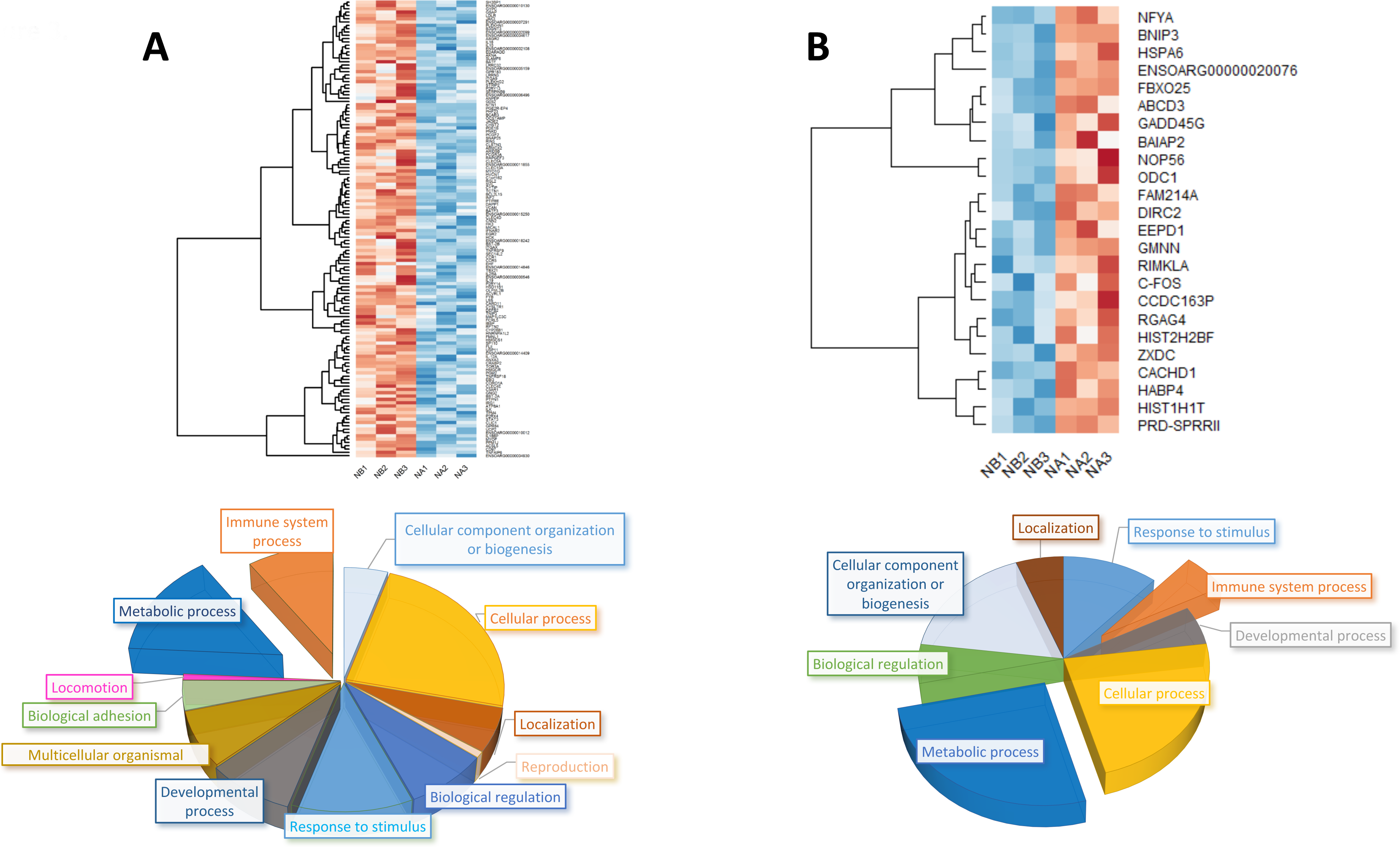
Differentialy expressed genes (DEG, padj < 0.1) in agonist (NA) compared to antagonist-preincubated LPS-exposed naïve microglia (NB). **A.** Heat map representation of differentially expressed down regulated genes and gene ontology pie chart. **B.** Heat map of up regulated genes in NA, compared to NB. Gene Ontology of each set of DEGs was presented as a pie chart at the bottom of each corresponding heat map. Note that “Immune system” and “Metabolic process” are both strongly down and up regulated, possibly referring to different functions being turned off while others are turned on under the influence of cholinergic signaling through α7nAChR.

Interestingly, we noticed that GO terms such as Locomotion (GO:0040011) and Reproduction (G0:0000003) were associated with DE down regulated genes (Figure 3A). We performed a second GO analysis with ToppGene of DEG down regulated, selected significant GO terms clusters (P < 10^−3^) and represented these in a bar chart with −Log(P) (Figure 4). Among DEG down regulated, the immune response was strongly significant (P = 6.10 × 10^−15^); we also noticed leukocyte migration and inflammatory response among the GO terms clustering (Figure 4, Table 3).

**Figure 4.**
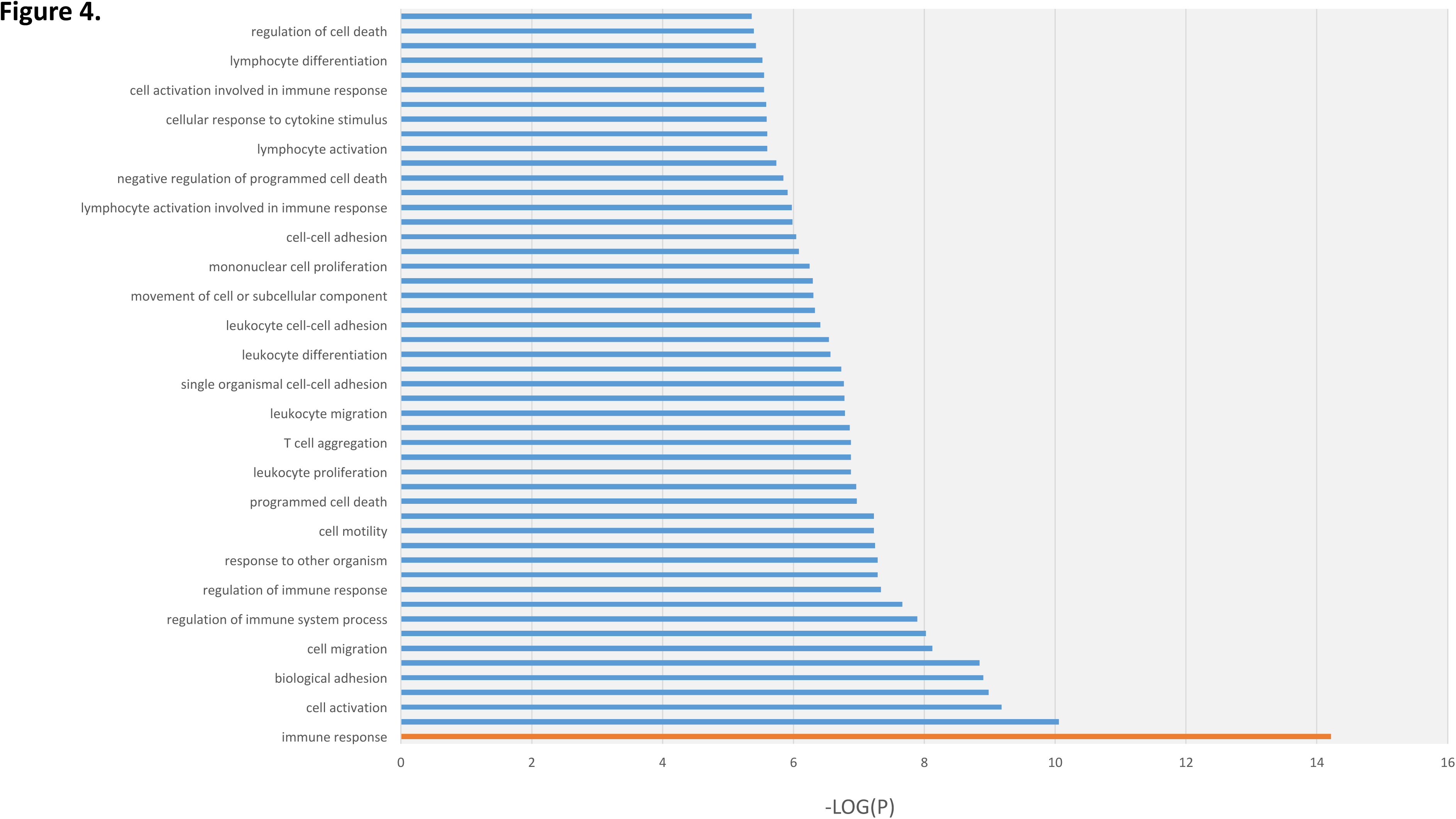
Gene ontology analysis with ToppGene of selected DEGs in agonist (NA) compared to antagonist-preincubated LPS-exposed naïve microglia (NB). Bar graph of 138 DEG down regulated. Each selected GO term (P < 10^−3^) was represented on a –Log (P) scale.

Lending support to our hypothesis, these results confirmed the anti-inflammatory effect of the α7nAChR agonistic stimulation on microglia.

#### Modulation of the memory of inflammation by α7nAChR signaling

From two differential analyses, NB versus NC and SHB versus NB, we selected up and down regulated genes with Log2 > |2| and represented common genes into a Venn diagram (Figure 5A). A total of 7 genes were DE and up regulated in both analyses, and two genes were down regulated. Interestingly, common down regulated genes were HMOX1 and SLC40A1.

**Figure 5.**
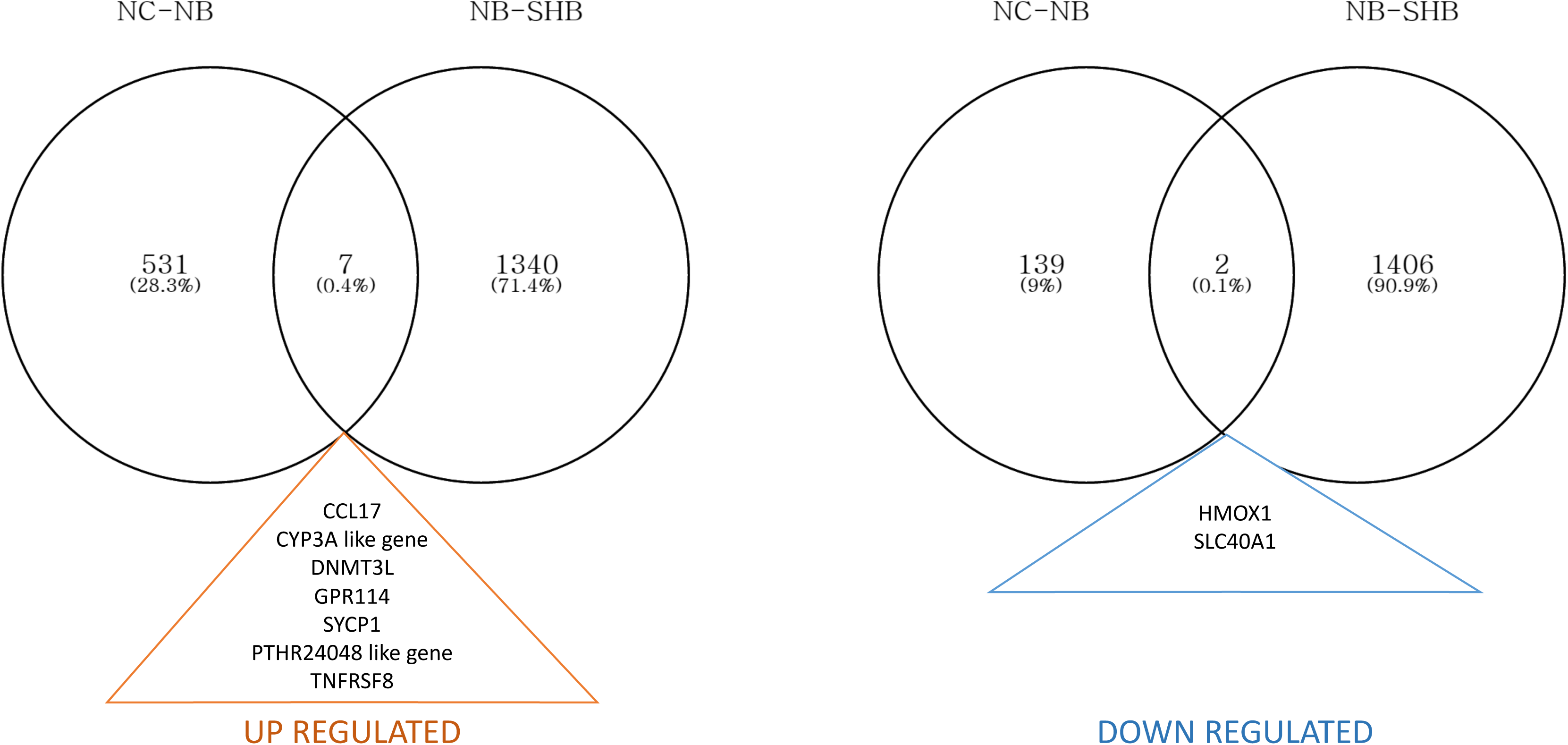
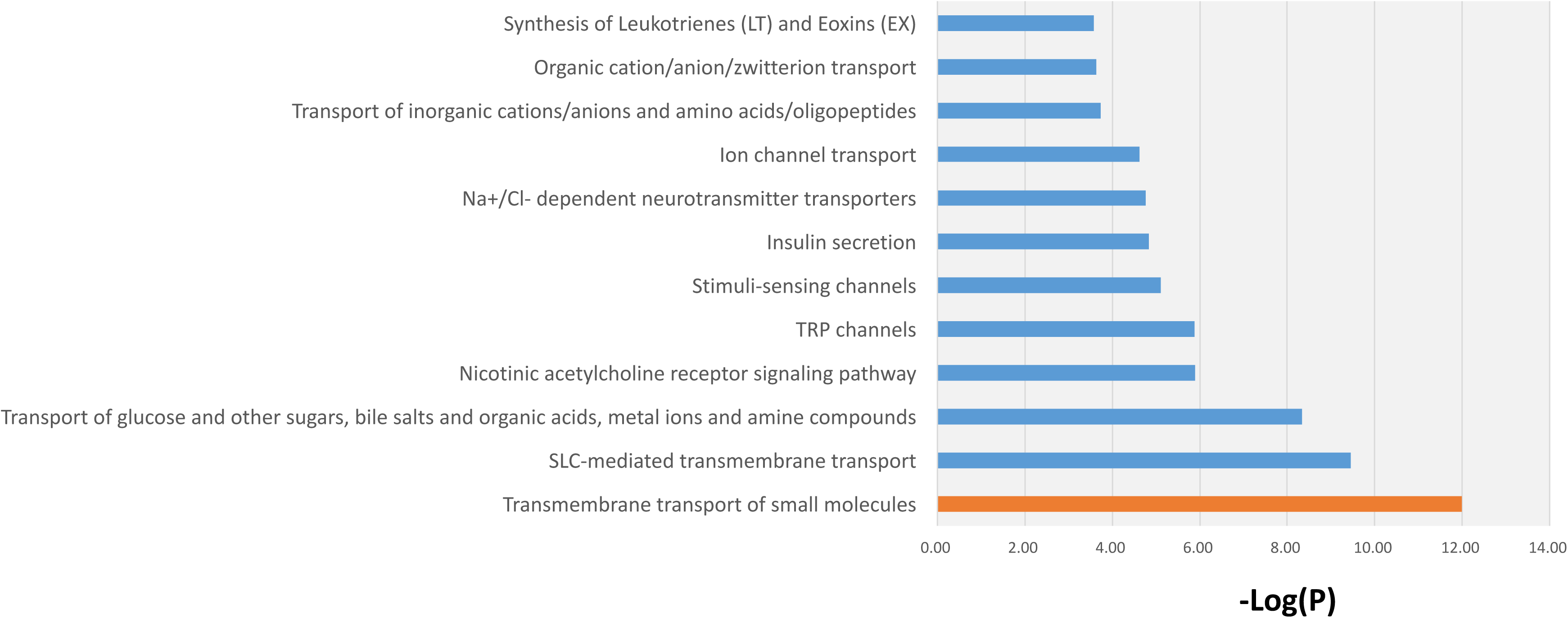
Stimulation with α-Bungarotoxin shows a unique transcriptome profile in microglia. **A.** Uniquely DEG in SHB (padj<0.01, Log2 > |2|). Each set of the common DEGs is written at the bottom of the Venn diagram. **B.** Pathways revealed by Gene Ontology of uniquely differentially expressed down regulated genes in SHB; pathways are represented with their −Log(P).

We reported the potential role of heme-oxigenase 1 (HMOX1) during neuroinflammation and will focus here on the gene SLC40A1 (ferroportin).(18) Using a double-hit model of LPS exposure, we showed that microglia gained memory of inflammation when pre-exposed to LPS *in vivo*, and we also pin-pointed the role of iron metabolism in this process. Here, gene ontology of uniquely down regulated genes in SHB versus NB contained, among pathways affected, metal ions compound (P = 4.64 × 10^−9^) and SLC-mediated transmembrane transport (P = 3.49 × 10^−10^). Similar to our earlier results with HMOX1 (18), this finding now highlights the putative role of SLC genes, another key component of the iron homeostasis in neuroinflammation, when cholinergic signaling is perturbed (Figure 5B).

Hepcidin (HAMP) plays a key role in linking inflammation and iron homeostasis (19), therefore we examined the RNA transcript level changes of hepcidin and ferroportin in our α7nAChR signaling model. Differential analyses of NL and NB to baseline and NA vs. NB revealed an opposite pattern of expression of HAMP and SLC40A1 in agonist-treated microglia, wherein ferroportin was up regulated (Log2 = 0.49) and hepcidin was down regulated (Log2 = - 1.31). However, only ferroportin was differentially expressed in naïve microglia and bungarotoxin-treated microglia (padj = 9.79 × 10^−4^, padj = 4.26 × 10^−5^, respectively). Further validation will be necessary to confirm these results (Figure 6).

**Figure 6.**
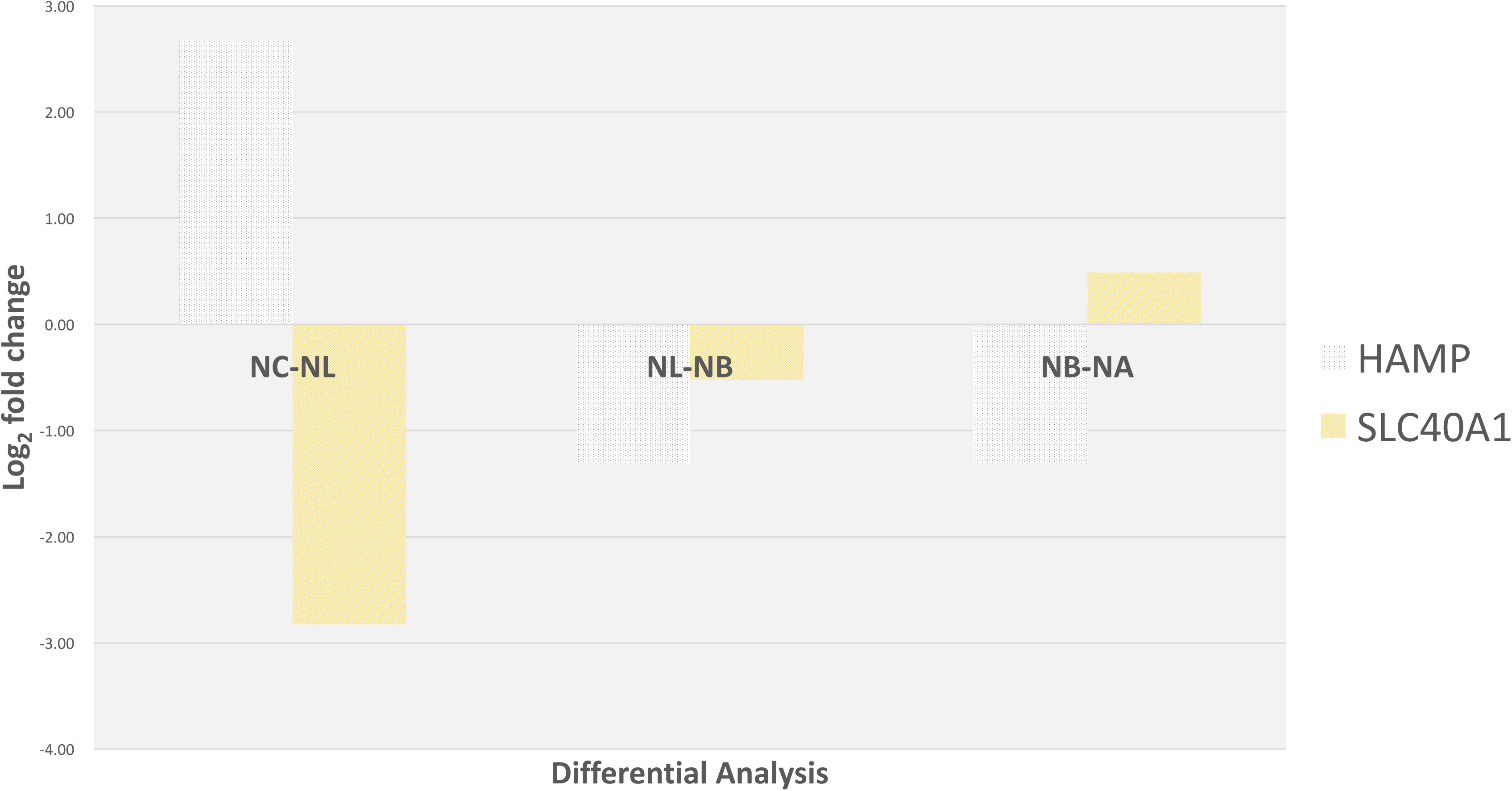
Genetic expression variation of SLC40A1 and HAMP through α7nAChR signaling. Each differential analysis is noted on the x-axis. The gene SLC40A1 is coding for the protein Ferroportin-1, a transmembrane protein, transporter of iron molecules into the bloodstream. Ferroportin-1 and HAMP have both an opposite pattern in agonist-treated microglia compared to naïve LPS-exposed microglia.

Similar to our findings on naïve microglia(18) and lending support to the epigenetic mechanisms of such neuroinflammation memory, here we also found HDAC1 up regulation following LPS exposure to be potentiated by pre-treatment with α7nAChR antagonist (NC-NB). This supports the notion that blocking α7nAChR signaling has pro-inflammatory effects not only on the level of cytokine secretion, but also on the level of strengthening the inflammation memory. HDAC6 behaved in the opposite direction of HDAC1, again similar to what we reported for LPS alone and potentiated by the α7nAChR antagonism (Table 3).

## DISCUSSION

In a unique double hit model of fetal neuroinflammation in a large mammalian brain, we studied transcriptomic changes following modulation of cholinergic signaling through α7nAChR with selective antagonistic drug α-Bungarotoxin and agonistic drug AR-R17779. We previously reported the activation of major inflammatory pathways JAK-STAT and NFKB in naïve microglia exposed to LPS as an inflammatory stimulus.(18) In our current analysis, we demonstrated the enhanced activation of these inflammatory pathways in microglia exposed to LPS and pre-treated with α-Bungarotoxin. Both, transcriptomic activation and protein level IL-1β secretion patterns were enhanced in α-Bungarotoxin pre-treated microglia compared to microglia exposed to LPS alone. Our findings extend the *in vitro* observations in mature rodent primary microglia cultures to a large mammalian developing brain exposed to LPS *in vivo* and *in vitro*.(15)

We aimed to understand biological processes in microglia when exposed to AR-R17779. The key finding from the differential analysis of NA versus NB was that genes of the immune response were DE down regulated in NA and most of these genes were up regulated in LPS-treated microglia (data not shown). Notably, JAK3 was DE down regulated in NA, and C-FOS was up regulated. DEG up regulated in NA and clustering in the immune system process GO term comprised only HSPA6 and GADD45G. The latter is known to regulate cytokine expression during LPS-induced inflammation (20), whereas HPSA6 is only induced under severe oxidative stress.(21) Lending support to our hypothesis, our findings in NA versus NB comparisons confirmed the anti-inflammatory effect of the α7nAChR agonistic stimulation in microglia.

The current findings represent the first direct *in vitro* validation of our earlier indirect *in vivo* observations in the fetal sheep brain of the same gestational age when we proposed the existence of a fetal cerebral cholinergic anti-inflammatory pathway with *in situ* evidence of the expression of α7nAChR on microglia.(22)

### Effect of α7nAChR signaling on the FBP / HMOX1 microglial phenotype

We reported HMOX1_down_ / FBP^up^ transcriptome phenotype in SHL microglia.(18) We expected that the agonistic stimulation of α7nAChR would attenuate this phenotype. HMOX1 and FBP were both differentially expressed in SHL microglia and this phenotype was sustained with antagonistic treatment. Differential expression in naïve microglia NA and NB did reveal an opposite expression, wherein HMOX1 was up regulated and FBP was down regulated in NA, however a statistically significant differential expression was not observed. Thus, further studies are needed to validate these results in the SHL microglia stimulated agonistically on the α7nAChR (cf. Methodological considerations).

### α7nAChR signaling modulates the epigenetic memory of inflammation in fetal microglia

Our findings suggest a potential role of histones in the memory of neuroinflammation.(18) Here, we asked whether epigenetic mechanisms are involved in enhancement and reduction of neuroinflammation in microglia via α7nAChR signaling. Differential analysis of NA versus NL revealed two DEG up regulated, HIST1H2BI and HIST1H1T, both corresponding to histone cluster 1H, strengthening our hypothesis of memory of neuroinflammation sustained by epigenetic factors(18) and extending it to involve α7nAChR signaling.

Our previous report showed that HDAC1, HDAC2 and HDAC6 were potentially involved in the memory of neuroinflammation. In this study, among selected HDAC genes DE in NB compared to NC, all showed an opposite expression pattern in NA compared to NB. The gene HAT1 was also DE down regulated in our previous report and was up regulated in the agonistically-treated microglia. Of note, similar to our published findings on naïve microglia, here we also found HDAC1 up regulation following LPS exposure and this effect was further potentiated by pre-treatment with α7nAChR antagonist (NC-NB) supporting the notion that blocking α7nAChR signaling has pro-inflammatory effects not only on the level of cytokine secretion, but also on the epigenetic level by strengthening the inflammation memory.

### Microglial iron homeostasis is modulated by α7nAChR signaling: implications for neurodevelopment

To further address mechanisms involved in memory of inflammation, we extracted common DE genes between naive and second hit microglia treated with α-Bungarotoxin. Common DEG up regulated included the genes CCL17, CYP3A and DNMT3L. CCL17 is known the mediate inflammation through macrophages (23), while CYP3A (cytochrome P450 3A) plays a major role in drug metabolism. CYP3A function in microglia is not known. Previous reports cited the up regulation of DNMT3L in TLR3- and TLR4 stimulated microglia (24), concordant with our results in α-Bungarotoxin treated microglia. We identified two DEG down regulated genes SLC40A1 and HMOX1. In conjunction with published results on HMOX1, we aimed to understand the role of SLC40A1, an iron-regulated transporter, known as ferroportin.

Our data indicate a combination of down regulation of metal ion transporter, ferroportin, with HMOX1. Ferroportin acts as a receptor for hepcidin (HAMP).(25) Hepcidin production is increased during the inflammatory response with increased binding of hepcidin to ferroportin leading to the internalization and degradation of ferroportin. This mechanism consequently suppresses enteral iron absorption and cellular iron release, whereas a decrease in hepcidin promotes iron uptake. Our results in fetal microglia are concordant with these published data in non-brain cells. Indeed, in naïve microglia (NL) compared to baseline (NC), HAMP was up regulated during inflammation and SLC40A1was down regulated. This expression pattern was reversed in α7nAChR agonist-treated (NA) compared to antagonist-treated microglia (NB). Our results suggest that in microglia, during inflammation, iron uptake is not only regulated by hepcidin binding to ferroportin but also by the level of transcription of ferroportin under cholinergic control.

Further studies are needed to clarify the role of hepcidin and heme-oxigenase during neuroinflammation and in memory of neuroinflammation. Such studies will help elucidate the mechanism underlying memory of neuroinflammation acquired *in utero* and the neurodevelopmental sequelae.

Evidence is emerging that iron overload is intricately involved in cognitive dysfunction with microglia priming or activation playing a key role in this process.(26-28) Excess intracellular iron may result from postoperative inflammation mediated by hepcidin or age-related iron accumulation under conditions of chronic inflammation. While most work in this area has been done in cultures or rat animal models, to our knowledge, this is the first report of inflammation-triggered changes in iron homeostasis in a larger mammalian brain with high resemblance to human physiology and in patterns of response to injury. We believe the current report is also the first observation of the putative link between cholinergic signaling in fetal microglia and the inflammatory milieu. Remarkably, iron homeostasis genes turn out to be key in determining the phenotype of the double-hit microglia. In light of the known role of iron in cognitive function (and dysfunction), our results raise the possibility that early disturbances in iron metabolism may have profound consequences in fetal and postnatal brain development. Our findings also suggest a possible therapeutic venue to modulate intracellular iron load via the α7nAChR.

### Impact of α7nAChR signaling manipulation on complement signaling pathway: putative implications for early programming of susceptibility to Alzheimer’s disease

Cognitive dysfunction may result not only from iron overload, but also from derangements in the neuronal-glial complement pathway interactions. Hyperactive microglia may prevent physiological synaptogenesis predisposing to Alzheimer’s disease (AD) later in life.(29),(30) The mediating pathway involves microglial-neuronal complement signaling suggesting that microglia could be potential early therapeutic targets in AD prevention or treatment and in other neurodegenerative diseases. The complement genes C1Q and CR3 (also known as CD11B) were involved in the microglia-mediated synaptic loss in a mouse model of AD.(29) Less is known about the function of the complement receptor 2 (CR2, also known as CD21) in neuroinflammation, especially in microglia. One study reports CR2^-/-^ mice to be more prone to neuronal injury with higher levels of astrocytosis following nerve root cord injury. This would suggest a neuroprotective role for CR2(31). Another study reports CR2^-/-^ mice subjected to traumatic brain injury to exhibit less astrocytosis and less microglial activation which would suggest the opposite role for CR2. (32) In mice, CR1 and CR2 are coded on the same gene and expressed as splice variants. In sheep, however, and other higher mammals these complement genes are coded separately. Hence, more studies are needed to gauge the functional role of CR2 in neuroinflammation, microglia in particular, especially with respect to human neurodegenerative diseases such as AD. Consequently, we conducted a secondary analysis of DEG in SHB compared to NB for CR2, C1Q chain A and B (C1QA, C1AB, respectively) and Complement component 3A Receptor 1 (C3AR1) as the best equivalent of CR3 in the annotated genome (Table 4). C3AR1 was the only gene showing a clear opposite pattern between agonistic and antagonistic α7nAChR treated microglia. Specifically, we found that agonistic stimulation of α7nAChR up regulated C3AR1 compared to antagonistic stimulation. We believe these findings deserve further study because pro-cholinergic drugs are used to treat AD symptoms, but the consequences of cholinergic stimulation on microglial signaling are not well understood. Our results suggest that blocking rather than enhancing α7nAChR signaling in microglia may be beneficial to help slow down synaptic degradation.

**Table 4.** α7nAChR signaling modulates the patterns of the complement C1Q-C3AR1 network activity implicated into microglial-neuronal interactions and pruning. Antagonistic stimulation of the microglial α7nAChR, but not the agonistic stimulation, down regulates both C1Q and C3AR1 expressions while up regulating C2. This is important to study further because pro-cholinergic drugs are used to treat AD symptoms, but it seems that at least in microglial α7nAChR signaling the opposite effect, blocking the cholinergic signaling, may be beneficial to help slow down synaptic degradation.

| Description | Gene | NC-NL | NB - SHB | NC - NA | NB - NA |
| --- | --- | --- | --- | --- | --- |
| Complement C1Q A chain | C1QA | -0.274 | **-2.379** | -0.081 | -0.227 |
| Complement C1Q B chain | C1QB | -0.093 | **-2.657** | 0.047 | -0.293 |
| Complement component 3a receptor 1 | C3AR1 | **1.662** | **-4.719** | 1.301 | -1.571 |
| Complement receptor type 2 | CR2 | 4.549 | **5.488** | 1.425 | 2.509 |

### Mechanistic model of interactions between iron homeostasis and α7nAChR signaling pathways in microglia

Based on our findings and the supporting literature, we propose a model of interactions between iron homeostasis and α7nAChR signaling in microglia (Fig. 7). The model highlights in red three exogenous factors that may be driving the microglial phenotype, some more intuitive (inflammation) than others (iron and stress). We briefly outline below the mechanistic connections to the “non-intuitive” factors and refer the interested reader to the cited references for details.

**Figure 7.**
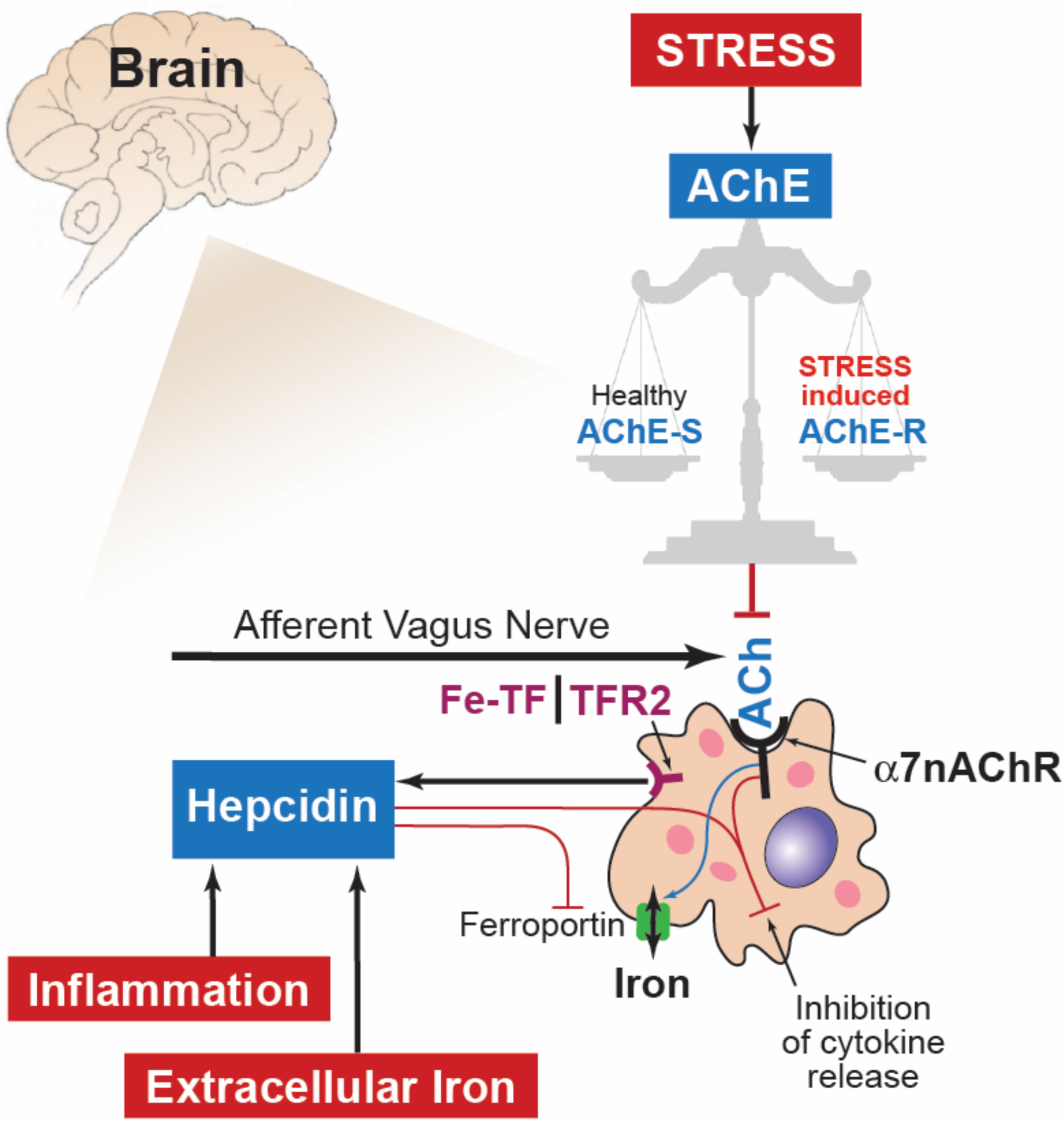
A model of interactions between iron homeostasis and α7nAChR signaling in microglia. Highlighted in red are the three exogenous factors that may be driving the microglial phenotype: inflammation, iron and stress. The former two stimulate hepcidin which in turn acts on ferroportin to be internalized and degraded. This reduces extracellular iron (sensed as Fe-TF, heme transferrin by TFR2, transferrin receptor 2 [shown here simplified as the representative iron sensor receptor] and increases the intracellular iron, a process referred to as iron sequestration. We propose that ferroportin’s membrane localization appears to also be controlled by the α7nAChR signaling (blue arrow). α7nAChR signaling depends on the acetyl choline (ACh) availability. The latter depends on the afferent vagus nerve cholinergic signaling in the brain via a distributed network as well as the non-vagal sources of ACh controlled by ACh esterase (AChE) activity and the availability of dietary choline. Remarkably, a large body of research has shown that AChE activity depends on chronic stress levels, a factor highly relevant in fetal microglia context in particular, because stress is very common in pregnancy. Stress results in shifts of the post-translational modification of AChE from AChE-S splice variant (healthy) to the less stable AChE-R variant.

Iron deficiency is the most common form of nutrient deficiency worldwide. According to the World Health Organization, it affects nearly 2 billion people and up to 50% of women who are pregnant.(33) At birth, 25-85% of premature babies are iron deficient and all will become iron deficient after birth, if not adequately supplemented, especially in developing countries.(34) Iron is essential for neonatal and long-term cognitive and physical development.(35),(36) Fetal or early postnatal inflammation may result in hepcidin-mediated intracellular microglial iron sequestration which may polarize microglia toward an inflammatory phenotype. We suggest a yet to be validated signaling pathway between the α7nAChR signaling cascade and the hepcidin-ferroportin steady-state. Such pathway may counteract to a degree hepcidin’s inhibitory effect on iron release due to the ferroportin internalization and degradation. We propose that such pathway is bidirectional with modulatory effects of hepcidin-ferroportin signaling on α7nAChR signaling. The bidirectional aspect is supported by our surprising observation of the IL-1ß secretion profile in the SHA group. *In utero* inflammation may reprogram microglial iron homeostasis toward iron sequestration which in turn diminishes the anti-inflammatory effect of the α7nAChR signaling. At second hit, the resulting net effect on the expression of pro-inflammatory cytokines controlled by NF-kB via the putative α7nAChR - ferroportin – hepcidin signaling network becomes pro-inflammatory which would explain the surprising switch of the IL-1β secretion profile in the SHA group compared to NA group. This is further supported by the finding that only accounting for *in utero* inflammation (first hit) revealed the SHB effect.

The second major factor is the actual availability of α7nAChR agonists to drive this pathway. One obvious candidate is the afferent cholinergic anti-inflammatory pathway via the vagus nerve that is comprised to ˜80% of afferent fibers and has wide-ranging projections in the brain.(37),(22),(38),(39) Another very well researched knob controlling ACh availability in the brain, independent of the vagal cholinergic signaling pathway, is the acetylcholinesterase (AChE). AChE is a key enzyme that regulates the ACh levels.(40) ACh binds to immune system cells such as microglia in the brain and macrophages in the periphery and decreases their propensity to respond to inflammatory stimuli during healthy (homeostatic) and infectious states (*e.g.*, bacterial sepsis).(41),(42) Under stress, elevated cortisol levels alter AChE gene expression to induce over-production of AChE and replace the major stable AChE splice variant AChE-S by the less stable AChE-R variant.(43, 44) Increased levels of AChE-R have been shown to result in chronic inflammation that ultimately impedes the body’s ability to defend itself against acute infections.(45, 46) AChE-S and –R ratios may influence ACh availability for the microglial α7nAChR signaling. The dependence on stress is particularly intriguing and relevant for neurodevelopment in the context of fetal microglial physiology, because stress during pregnancy is a very common phenomenon with estimates ranging from conservative 10% to as high as 50% of all pregnant women to report at least one major stress event during pregnancy.(47),(48, 49)

### Methodological considerations

Aside from the lack of biological replicates for SHL microglia, we conducted differential analyses in all other samples. The lack of SHA replicates for sequencing prevented us from studying directly the effect of agonistic drugs on SHL microglia. Thus, further study is needed on the transcriptomic level in α7nAChR agonistic stimulated SHL microglia.

## METHODS

### In vitro microglia culture protocol

Fetal sheep brain tissues were obtained during sheep autopsy after completion of the experiment for *in vitro* study. The non-instrumented, untreated twins were designated “naïve” (no LPS exposure *in vivo).* Instrumented animals that received LPS *in vivo* were used for 2^nd^ hit LPS exposure *in vitro.* Fetal sheep microglia culture protocol was adapted from an established human adult and fetal microglia culture protocol that was modified to include a myelin removal step following high-speed centrifugation.(50) Briefly, fetal sheep cells were plated on poly-L-lysine (PLL)-coated tissue culture flasks at a concentration of 2 × 10^6^ cells /ml in DMEM with 5% heat-inactivated fetal bovine serum (Gibco, Canada Origin), 1% penicillin/ streptomycin, and 1% glutamine (5% DMEM), in which microglia are preferable to grow.(50) Cells were allowed to incubate for seven days at 37°C, 5% CO_2_, followed by media change by centrifugation and addition of re-suspended cells back to the culture flask. Cells were continued to incubate for seven more days with 5% DMEM at 37°C, 5% CO_2_, before floating cells were collected. Carefully collecting the floating microglia to avoid contamination with astrocytes and oligodendrocytes, the cells were incubated in 24-well plate at 1 × 10^5^ cells/mL with 5% DMEM for another 4-5 days, and then treated with or w/o LPS (100ng/ml, Sigma L5024, from E coli O127, B8)) for 6h. Cell conditioned media were collected for cytokine analysis, 0.5ml TriZol were added per well for RNA extraction. To verify microglia purity, a portion of floating cells were cultured in 24-well plate at above conditions for flow cytometry analysis (see below), cell morphology was documented with light microscopy. Another portion of floating cells were plated into Lab-Tek 8 well chamber glass slide (Thermo Scientific) and treated with or w/o LPS for immunocytochemistry analysis, in this experiment, some wells of astrocytes cultured at DMEM with 10% FCS were included for comparison.

#### Cell culture

Microglia cells isolation and culture were described in detail elsewhere.(18) Briefly, prior to exposure to LPS, cells were pre-treated for 1 hour with either 10nM AR-R17779 hydrochloride (Tocris Bioscience Cat# 3964), a selective α7nAChR agonist, or 100nM α-Bungarotoxin (Tocris Bioscience Cat# 2133), a selective α7nAChR antagonist. Optimal dose of AR-R17779 (A) or α-Bungarotoxin (B) was chosen based on a dose-response experiment with LPS; AR-R17779 hydrochloride was dissolved in DMSO into a stock solution. We then made serial dilutions, whereas α-Bungarotoxin was reconstituted with culture media into a stock solution and underwent serial dilutions. AR-R17779 and α-Bungarotoxin preparations were added well by well; the same volume of vehicle (either DMSO or cell culture media) was added in control wells. Therefore, in a complete cell culture experiment, we had four experimental groups: Control, LPS, LPS+B and LPS +A. Second hit cell cultures were designed with the same pattern and divided into four experimental groups: Control (SHC), LPS (SHL), LPS+B (SHB) and LPS+A (SHA).

#### Measurements of inflammatory responses

##### Measurement of cytokines in plasma and cell culture media

Cytokine concentrations in cell culture media (IL-1β) were determined by using an ovine-specific sandwich ELISA. Briefly, 96-well plates (Nunc Maxisorp, high capacity microtitre wells) were pre-coated with the capture antibody, the mouse anti sheep monoclonal antibodies (IL-1β, MCA1658, Bio Rad AbD Serotec) at a concentration 4μg/ml on ELISA plate at 4°C for overnight, after 3 times wash with washing buffer (0.05% Tween 20 in PBS, PBST), plates were then blocked for 1h with 1% BSA in PBST for plasma samples or 10% FBS for cell culture media. Recombinant sheep proteins (IL-1 β, Protein Express Cat. no 968-405) were used as ELISA standard. All standards and samples (50μ! per well) were run in duplicate. Rabbit anti sheep polyclonal antibodies (detection antibody IL-1ß, AHP423, Bio Rad AbD Serotec) at a concentration of 4μg/ml were applied in wells and incubated for 30min at room temperature. Plates were washed with washing buffer for 5-7 times between each step. Detection was accomplished by assessing the conjugated enzyme activity (goat anti-rabbit IgG-HRP, dilution 1:5000, Jackson ImmunoResearch, Cat. No 111-035-144) via incubation with TMB substrate solution (BD OptEIA TMB substrate Reagent Set, BD Biosciences Cat. No 555214), colour development reaction was stopped with 2N sulphuric acid. Plates were read on ELISA plate reader at 450nm, with 570nm wavelength correction (EnVision 2104 Multilabel Reader, Perkin Elmer). The sensitivity of IL-1ß ELISA for media was 41.3 pg/ml. For all assays, the intra-assay and inter-assay coefficients of variance was <5% and <10%, respectively.

##### RNAseq approach

The overall experimental design was divided into three phases: sequencing, quantification and discovery (Figure 1A). RNA extraction and RNA quantification: Total RNA was extracted from cultured microglia using TRIzol Reagent (Life Technologies), to obtain enough amount RNA, same treatment cells maybe pooled in same replicates. RNA quantity and quality (RNA integrity number, RIN) was established by using a RNA Nano Chips (Agilent RNA 6000 Nano Chips) with Agilent 2100 BioAnalyzer. All samples had acceptable RIN value ranging from 6 to 8.5 A total of 12 naïve microglia cultures from four sets of replicates were selected for RNA sequencing at high throughput, as well as three second hit microglia cultures, including SHC, SHL and antagonist-exposed microglia (SHB). Second hit microglia further exposed to agonistic drugs (SHA) were not sequenced due to low RIN. This is left for future studies.

RNAseq libraries were prepared by using Illumina TruSeq RNA Sample Preparation v2 kit (Illumina) and quality control was performed on the BioAnalyzer. Single-end 50-bp sequencing was performed at high throughput on an Illumina HiSeq2500 at the CHU Ste-Justine Core Facility Sequencing Platform.

##### RNAseq data analysis

###### Reads alignment to the reference genome

To maximize the number of genes covered, raw data were mapped to the reference genome of the sheep *Ovis aris* v3.1 from NCBI and Ensembl (GCA_000298735.1) as transcriptome reference. Index of the reference fasta file were built with Bowtie2(51), we then trimmed the adaptor of the fastQ files with TrimGalore, and mapped reads to the reference with Tophat2.(52) From the aligned reads from Tophat2, the number of reads per gene were counted with HTseq and assembled into a matrix containing the read count of each gene per sample.(53)

###### Normalization and transcriptome analysis

In order to find differentially expressed genes we used DESeq2 to normalize the dataset, generate Log2-fold changes and adjacent P values (padj).(54) We performed 6 differential analyses of microglial transcriptome (Table 1). After stimulation through α7nAChR with agonistic and antagonistic drugs, we aimed to measure changes at the transcriptome level. Thus, we eliminated the background carried by exposure to LPS by comparing directly NA read count variation to NB. Due to the lack of replicates in our second-hit experiment, we were unable to perform thorough differential analysis of second hit microglial cells and leave the second hit transcriptome analysis for future studies. A gene was considered differentially expressed if its adjacent p-value was strictly lower than 0.1. Pools of up and down regulated genes and differentially expressed genes were clustered and visualized into heat maps, generated in R using the log2 normalized counts and the heatmap.2 method of the gplots library.(55)

###### Gene selection and Gene Ontology (GO)

The sheep genome is not yet supported by most gene ontology platforms; therefore, downstream analyses were performed with orthologs in the human genome *Homo sapiens.* To select relevant genes among up regulated and down regulated genes, we performed gene enrichment analysis for biological process and molecular function with ToppGenes and FDR < 0.05.(56, 57) Bar diagram of significant GO terms (P < 10^−3^) was presented on a –Log (P) scale. Protein-protein interaction networks were generated with the STRING database and disconnected nodes were not represented.(58) Gene Ontology was also performed in parallel with PantherDB and only biological processes were presented in pie charts.(59)

###### STATISTICS

Generalized estimating equations (GEE) modeling approach was used to assess the effects of LPS and drug treatments. We used a linear scale response model with LPS/drug treatment group and presence or absence of second hit exposure as predicting factors to assess their interactions using maximum likelihood estimate and Type III analysis with Wald Chi-square statistics. SPSS Version 21 was used for these analyses (IBM SPSS Statistics, IBM Corporation, Armonk, NY). Significance was assumed for p < 0.05. Results are provided as means ± SEM or as median {25-75} percentile, as appropriate. Not all measurements were obtained for each animal studied.

###### Study approval

This study was carried out in strict accordance with the recommendations in the Guide for the Care and Use of Laboratory Animals of the National Institutes of Health. The respective *in vivo* and *in vitro* protocols were approved by the Committee on the Ethics of Animal Experiments of the Université de Montréal (Permit Number: 10-Rech-1560).

## AUTHOR CONTRIBUTIONS

CSM, PB, GF, AD, JPA and MGF are responsible for the conception and design.

MCa, HLL, CSM, LDD, PB, GF, and AD acquired data.

MCo, MCa, HLL, CSM, LBB, JPA and MGF did the analysis and interpretation of data.

MCo, MCa and MGF drafted the manuscript.

MCo, MCa and MGF are responsible for revising it for intellectual content.

MCo, MCa, HLL, CSM, LDD, PB, GF, AD, LBB, JPA and MGF gave final approval of the completed manuscript.

## ACKNOWLEDGEMENTS

The authors thank Dora Siontas, Manon Blain for cell culture and ICC, Lamia Naouel Hachehouche for cytokines ELISA assay, Vania Yotova for RNAseq library preparation, Jean-Christopher Grenier for alignment to the reference genome and read count, St-Hyacinthe CHUV team and M. Michel-Robinson for technical assistance and Jan Hamanishi for graphical design.

## References

1. Saigal S, and Doyle LW. An overview of mortality and sequelae of preterm birth from infancy to adulthood. Lancet. 2008;371(9608):261–9.

2. Rees S, and Inder T. Fetal and neonatal origins of altered brain development. Early human development. 2005;81(9):753–61.

3. Murthy V, and Kennea NL. Antenatal infection/inflammation and fetal tissue injury. Best practice & researchClinical obstetrics & gynaecology. 2007;21(3):479–89.

4. Polin RA. Systemic infection and brain injury in the preterm infant. Jornal de pediatria. 2008;84(3):188–91.

5. Hagberg H, Peebles D, and Mallard C. Models of white matter injury: comparison of infectious, hypoxic-ischemic, and excitotoxic insults. Mental retardation and developmental disabilities research reviews. 2002;8(1):30–8.

6. Wang X, Rousset CI, Hagberg H, and Mallard C. Lipopolysaccharide-induced inflammation and perinatal brain injury. Seminars in fetal & neonatal medicine. 2006;11(5):343–53.

7. Gotsch F, Romero R, Kusanovic JP, Mazaki-Tovi S, Pineles BL, Erez O, Espinoza J, and Hassan SS. The fetal inflammatory response syndrome. Clinical obstetrics and gynecology. 2007;50(3):652–83.

8. Fahey JO. Clinical management of intra-amniotic infection and chorioamnionitis: a review of the literature. Journal of midwifery & women's health. 2008;53(3):227–35.

9. Fishman SG, and Gelber SE. Evidence for the clinical management of chorioamnionitis. Semin Fetal Neonatal Med. 2012;17(1):46–50.

10. Agrawal V, and Hirsch E. Intrauterine infection and preterm labor. Semin Fetal Neonatal Med. 2012;17(1):12–9.

11. Karrow NA. Activation of the hypothalamic-pituitary-adrenal axis and autonomic nervous system during inflammation and altered programming of the neuroendocrine-immune axis during fetal and neonatal development: lessons learned from the model inflammagen, lipopolysaccharide. Brain, behavior, and immunity. 2006;20(2):144–58.

12. Bilbo SD, and Schwarz JM. Early-life programming of later-life brain and behavior: a critical role for the immune system. Frontiers in behavioral neuroscience. 2009;3(14.

13. van der Valk P, and Amor S. Preactive lesions in multiple sclerosis. Current opinion in neurology. 2009;22(3):207–13.

14. Kettenmann H, Hanisch UK, Noda M, and Verkhratsky A. Physiology of microglia. Physiological reviews. 2011;91(2):461–553.

15. Shytle RD, Mori T, Townsend K, Vendrame M, Sun N, Zeng J, Ehrhart J, Silver AA, Sanberg PR, and Tan J. Cholinergic modulation of microglial activation by alpha 7 nicotinic receptors. Journal of neurochemistry. 2004;89(2):337–43.

16. Suzuki T, Hide I, Matsubara A, Hama C, Harada K, Miyano K, Andra M, Matsubayashi H, Sakai N, Kohsaka S, et al. Microglial αlpha7 nicotinic acetylcholine receptors drive a phospholipase C/IP3 pathway and modulate the cell activation toward a neuroprotective role. JNeurosci Res. 2006;83(8):1461–70.

17. Hua S, Ek CJ, Mallard C, and Johansson ME. Perinatal hypoxia-ischemia reduces alpha 7 nicotinic receptor expression and selective alpha 7 nicotinic receptor stimulation suppresses inflammation and promotes microglial Mox phenotype. Biomed Res Int. 2014;2014(718769.

18. Cao M, Cortes M, Moore CS, Leong SY, Durosier LD, Burns P, Fecteau G, Desrochers A, Auer RN, Barreiro LB, et al. Fetal microglial phenotype in vitro carries memory of prior in vivo exposure to inflammation. Front Cell Neurosci. 2015;9(294).

19. Robb A, and Wessling-Resnick M. Regulation of transferrin receptor 2 protein levels by transferrin. Blood. 2004;104(13):4294–9.

20. Schmitz I. Gadd45 proteins in immunity. Adv Exp Med Biol. 2013;793(51-68.

21. Ito YA, Goping IS, Berry F, and Walter MA. Dysfunction of the stress-responsive FOXC1 transcription factor contributes to the earlier-onset glaucoma observed in Axenfeld-Rieger syndrome patients. Cell Death Dis. 2014;5(e1069.

22. Frasch MG, Szynkaruk M, Prout AP, Nygard K, Cao M, Veldhuizen R, Hammond R, and Richardson BS. Decreased neuroinflammation correlates to higher vagus nerve activity fluctuations in near-term ovine fetuses: a case for the afferent cholinergic anti-inflammatory pathway? Journal of neuroinflammation. 2016;13(1):103.

23. Achuthan A, Cook AD, Lee MC, Saleh R, Khiew HW, Chang MW, Louis C, Fleetwood AJ, Lacey DC, Christensen AD, et al. Granulocyte macrophage colony-stimulating factor induces CCL17 production via IRF4 to mediate inflammation. J Clin Invest. 2016;126(9):3453–66.

24. Das A, Chai JC, Kim SH, Lee YS, Park KS, Jung KH, and Chai YG. Transcriptome sequencing of microglial cells stimulated with TLR3 and TLR4 ligands. BMC Genomics. 2015;16(517.

25. Wessling-Resnick M. Iron imports. III. Transfer of iron from the mucosa into circulation. Am J Physiol Gastrointest Liver Physiol. 2006;290(1):G1–6.

26. Li Y, Pan K, Chen L, Ning JL, Li X, Yang T, Terrando N, Gu J, and Tao G. Deferoxamine regulates neuroinflammation and iron homeostasis in a mouse model of postoperative cognitive dysfunction. Journal of neuroinflammation. 2016;13(1):268.

27. Ward RJ, Zucca FA, Duyn JH, Crichton RR, and Zecca L. The role of iron in brain ageing and neurodegenerative disorders. Lancet Neurol. 2014;13(10):1045–60.

28. Urrutia P, Aguirre P, Esparza A, Tapia V, Mena NP, Arredondo M, Gonzalez-Billault C, and Nunez MT. Inflammation alters the expression of DMT1, FPN1 and hepcidin, and it causes iron accumulation in central nervous system cells. JNeurochem. 2013;126(4):541–9.

29. Hong S, Beja-Glasser VF, Nfonoyim BM, Frouin A, Li S, Ramakrishnan S, Merry KM, Shi Q, Rosenthal A, Barres BA, et al. Complement and microglia mediate early synapse loss in Alzheimer mouse models. Science. 2016;352(6286):712–6.

30. Stephan AH, Barres BA, and Stevens B. The complement system: an unexpected role in synaptic pruning during development and disease. Annu Rev Neurosci. 2012;35(369-89.

31. Lindblom RP, Berg A, Strom M, Aeinehband S, Dominguez CA, Al Nimer F, Abdelmagid N, Heinig M, Zelano J, Harnesk K, et al. Complement receptor 2 is up regulated in the spinal cord following nerve root injury and modulates the spinal cord response. J Neuroinflammation. 2015;12(192.

32. Neher MD, Rich MC, Keene CN, Weckbach S, Bolden AL, Losacco JT, Patane J, Flierl MA, Kulik L, Holers VM, et al. Deficiency of complement receptors CR2/CR1 in Cr2(-)/(-) mice reduces the extent of secondary brain damage after closed head injury. J Neuroinflammation. 2014;11(95.

33. Radlowski EC, and Johnson RW. Perinatal iron deficiency and neurocognitive development. Frontiers in human neuroscience. 2013;7(585.

34. Rao R, and Georgieff MK. Iron therapy for preterm infants. Clin Perinatol. 2009;36(1):27–42.

35. Lieblein-Boff JC, McKim DB, Shea DT, Wei P, Deng Z, Sawicki C, Quan N, Bilbo SD, Bailey MT, McTigue DM, et al. Neonatal E. coli infection causes neuro-behavioral deficits associated with hypomyelination and neuronal sequestration of iron. JNeurosci. 2013;33(41):16334–45.

36. Doom JR, and Georgieff MK. Striking while the iron is hot: Understanding the biological and neurodevelopmental effects of iron deficiency to optimize intervention in early childhood. Curr Pediatr Rep. 2014;2(4):291–8.

37. Hosoi T, Okuma Y, and Nomura Y. Electrical stimulation of afferent vagus nerve induces IL-1beta expression in the brain and activates HPA axis. Am J Physiol Regul Integr Comp Physiol. 2000;279(1):R141–7.

38. Fraschini M, Demuru M, Puligheddu M, Floridia S, Polizzi L, Maleci A, Bortolato M, Hillebrand A, and Marrosu F. The re-organization of functional brain networks in pharmaco-resistant epileptic patients who respond to VNS. Neurosci Lett. 2014;580(153-7.

39. Kwan H, Garzoni L, Liu HL, Cao M, Desrochers A, Fecteau G, Burns P, and Frasch MG. VAGUS NERVE STIMULATION FOR TREATMENT OF INFLAMMATION:SYSTEMATIC REVIEW OF ANIMAL MODELS AND CLINICAL STUDIES. Bioelectronic Medicine. 2016;3(1-6.

40. Soreq H, and Seidman S. Acetylcholinesterase--new roles for an old actor. Nat Rev Neurosci. 2001;2(4):294–302.

41. Andersson U, and Tracey KJ. Reflex principles of immunological homeostasis. Annu Rev Immunol. 2012;30(313-35.

42. Soreq H. Checks and balances on cholinergic signaling in brain and body function. Trends Neurosci. 2015;38(7):448–58.

43. Friedman A, Kaufer D, Shemer J, Hendler I, Soreq H, and Tur-Kaspa I. Pyridostigmine brain penetration under stress enhances neuronal excitability and induces early immediate transcriptional response. Nat Med. 1996;2(12):1382–5.

44. Kaufer D, Friedman A, Seidman S, and Soreq H. Acute stress facilitates long-lasting changes in cholinergic gene expression. Nature. 1998;393(6683):373–7.

45. Sternfeld M, Shoham S, Klein O, Flores-Flores C, Evron T, Idelson GH, Kitsberg D, Patrick JW, and Soreq H. Excess "read-through" acetylcholinesterase attenuates but the "synaptic" variant intensifies neurodeterioration correlates. Proc Natl Acad Sci U S A. 2000;97(15):8647–52.

46. Shaked I, Meerson A, Wolf Y, Avni R, Greenberg D, Gilboa-Geffen A, and Soreq H. MicroRNA-132 potentiates cholinergic anti-inflammatory signaling by targeting acetylcholinesterase. Immunity. 2009;31(6):965–73.

47. Lee AM, Lam SK, Sze Mun Lau SM, Chong CS, Chui HW, and Fong DY. Prevalence, course, and risk factors for antenatal anxiety and depression. Obstet Gynecol. 2007;110(5):1102–12.

48. Robinson AM, Benzies KM, Cairns SL, Fung T, and Tough SC. Who is distressed? A comparison of psychosocial stress in pregnancy across seven ethnicities. BMC pregnancy and childbirth. 2016;16(1):215.

49. Biaggi A, Conroy S, Pawlby S, and Pariante CM. Identifying the women at risk of antenatal anxiety and depression: A systematic review. J Affect Disord. 2016;191(62-77.

50. Durafourt BA, Moore CS, Blain M, and Antel JP. Isolating, culturing, and polarizing primary human adult and fetal microglia. Methods Mol Biol. 2013;1041(199-211.

51. Langmead B, and Salzberg SL. Fast gapped-read alignment with Bowtie 2. Nat Methods. 2012;9(4):357–9.

52. Kim D, Pertea G, Trapnell C, Pimentel H, Kelley R, and Salzberg SL. TopHat2: accurate alignment of transcriptomes in the presence of insertions, deletions and gene fusions. Genome Biol. 2013;14(4):R36.

53. Anders S, Pyl TP, and Huber W. HTSeq — A Python framework to work with high-throughput sequencing data. bioRxiv. 2014.

54. Love MI, Huber W, and Anders S. Moderated estimation of fold change and dispersion for RNA-Seq data with DESeq2. bioRxiv. 2014.

55. Warnes GR. 2008.

56. Chen J, Bardes EE, Aronow BJ, and Jegga AG. ToppGene Suite for gene list enrichment analysis and candidate gene prioritization. Nucleic Acids Res. 2009;37(Web Server issue):W305–11.

57. Kaimal V, Bardes EE, Tabar SC, Jegga AG, and Aronow BJ. ToppCluster: a multiple gene list feature analyzer for comparative enrichment clustering and network-based dissection of biological systems. Nucleic Acids Res. 2010;38(Web Server issue):W96–102.

58. Franceschini A, Szklarczyk D, Frankild S, Kuhn M, Simonovic M, Roth A, Lin J, Minguez P, Bork P, von Mering C, et al. STRING v9.1: protein-protein interaction networks, with increased coverage and integration. Nucleic Acids Res. 2013;41(Database issue):D808–15.

59. Mi H, Poudel S, Muruganujan A, Casagrande JT, and Thomas PD. PANTHER version 10: expanded protein families and functions, and analysis tools. Nucleic Acids Res. 2016;44(D1):D336–42.

